# Comprehensive Immune Monitoring of Clinical Trials to Advance Human Immunotherapy

**DOI:** 10.1101/489765

**Authors:** Felix J. Hartmann, Joel Babdor, Pier Federico Gherardini, El-Ad D. Amir, Kyle Jones, Bita Sahaf, Diana M. Marquez, Peter Krutzik, Erika O’Donnell, Natalia Sigal, Holden T. Maecker, Everett Meyer, Matthew H. Spitzer, Sean C. Bendall

## Abstract

The success of immunotherapy has led to a myriad of new clinical trials. Connected to these trials are efforts to discover biomarkers providing mechanistic insight and predictive signatures for personalization. Still, the plethora of immune monitoring technologies can face investigator bias, missing unanticipated cellular responses in limited clinical material. We here present a mass cytometry workflow for standardized, systems-level biomarker discovery in immunotherapy trials. To broadly enumerate human immune cell identity and activity, we established and extensively assessed a reference panel of 33 antibodies to cover major cell subsets, simultaneously quantifying activation and immune checkpoint molecules in a single assay. The resulting assay enumerated ≥ 98% of peripheral immune cells with ≥ 4 positively identifying antigens. Robustness and reproducibility were demonstrated on multiple samples types, across research centers and by orthogonal measurements. Using automated analysis, we monitored complex immune dynamics, identifying signatures in bone-marrow transplantation associated graft-versus-host disease. This validated and available workflow ensures comprehensive immunophenotypic analysis, data comparability and will accelerate biomarker discovery in immunomodulatory therapeutics.

## Introduction

Treating cancer via modulation of the immune system has recently shown curative clinical benefit in multiple types of cancer for which conventional chemotherapy has not worked. Three of the most widely employed strategies are hematopoietic stem cell transplantation, immune checkpoint blockade (Ribas and Wolchok, 2018) and adoptive transfer of chimeric antigen receptor (CAR) T cells (June et al., 2018), although many other approaches are being developed. To further investigate the immunotherapeutic potential of all approaches and combinations thereof, thousands of clinical trials are currently being planned and conducted (Farkona et al., 2016).

Many immunotherapy trials are accompanied by immune monitoring, which can provide crucial insights into immune cell behavior at both population and single-cell levels. Comprehensive phenotyping of immune populations aids in the elucidation of the cellular mechanisms underlying newly developed therapeutic approaches. It can also identify the presence of cellular and molecular signatures that stratify patients into distinct risk groups and/or help to predict clinical responses to therapy. The tremendous complexity and heterogeneity of the human immune system necessitates the use of single-cell technologies for its analysis. While flow cytometry has traditionally been the mainstay for such immune-monitoring applications, the advent of mass cytometry (i.e. cytometry by time-of-flight; CyTOF) now provides an opportunity to simultaneously quantify more molecular features while reducing signal overlap and background noise (Bandura et al., 2009; Bendall et al., 2011). The high-dimensional capabilities of mass cytometry enable the identification of a wide array of immune populations and cellular states in a single assay which allows comprehensive immune monitoring of small sample quantities but also across millions of cells from large groups of patients (Spitzer and Nolan, 2016).

Prior to using mass cytometry for immune cell phenotyping in clinical trials, rigorous validation studies must be performed to establish proper experimental and analytical workflows. While such specialized workflows have been developed independently at some research institutions, published studies using these methods are typically not comparable since each workflow uses distinct antibody panels to identify different target immune cell populations. Furthermore, the scope of a given study is often limited to specific target populations hypothesized to be of importance instead of broadly surveying all immune cell subsets in a given sample. This approach likely biases the analysis and overlooks unanticipated, potentially novel effects on other immune cell populations. Further, the unbiased analysis of such studies requires researchers to establish dedicated analytical frameworks to more effectively mine the high-dimensional datasets generated by mass cytometry (Arvaniti and Claassen, 2017; Bruggner et al., 2014; Nowicka et al., 2017).

To address these issues, we here present a mass cytometry-based experimental workflow for comprehensive, immune monitoring of cancer immunotherapy clinical trials. The proposed reference antibody panel used in this workflow is comprised of a readily available, established and validated set of 33 surface and intracellular antibodies, enabling the robust identification of key immune cell populations and cell states in a single assay. We achieved assignment of 98% of peripheral immune cells by positivity of four or more antigens. Importantly, the design facilitates the space for additional (≥10) targets without disruption of the core reference panel in order to address experiment-specific hypotheses, providing an unprecedented level of flexibility and customization compared to other workflows. Exemplifying this ability, we identify additional B cell maturation states and characterize myeloid cell heterogeneity across matched primary tumors and lymph node metastases, suggesting tissue-dependent expression of co-stimulatory molecules (CD86). Finally, we demonstrate the utility of this framework by monitoring immune cell reconstitution and identifying disease-associated immune signatures using an automated pipeline following bone marrow transplantation (BMT) in leukemia patients (n = 15). Together, this workflow provides a standardized immune monitoring approach that can greatly improve understanding of key molecular and cellular factors that influence and can predict therapeutic success and failure, providing biomarkers to improve the application of next generation treatments.

## Results

### Comprehensive phenotyping for human immunotherapy trials

In order to build a comprehensive human immunophenotyping pael for a single-pass analysis, we took a cell-lineage agnostic approach in order to maximize coverage of all immune populations expected in biological specimens (i.e. peripheral blood and tissue) from immunotherapy trials. As such, we first selected the major immune cell lineages and their subsets that would be ideal to detect in human cancer samples. This list is comprised of T cells, B cells, natural killer (NK) cells and various myeloid and granulocyte populations, thus covering all major immune cell lineages typically found (Fig. 1A).

**Figure 1.**
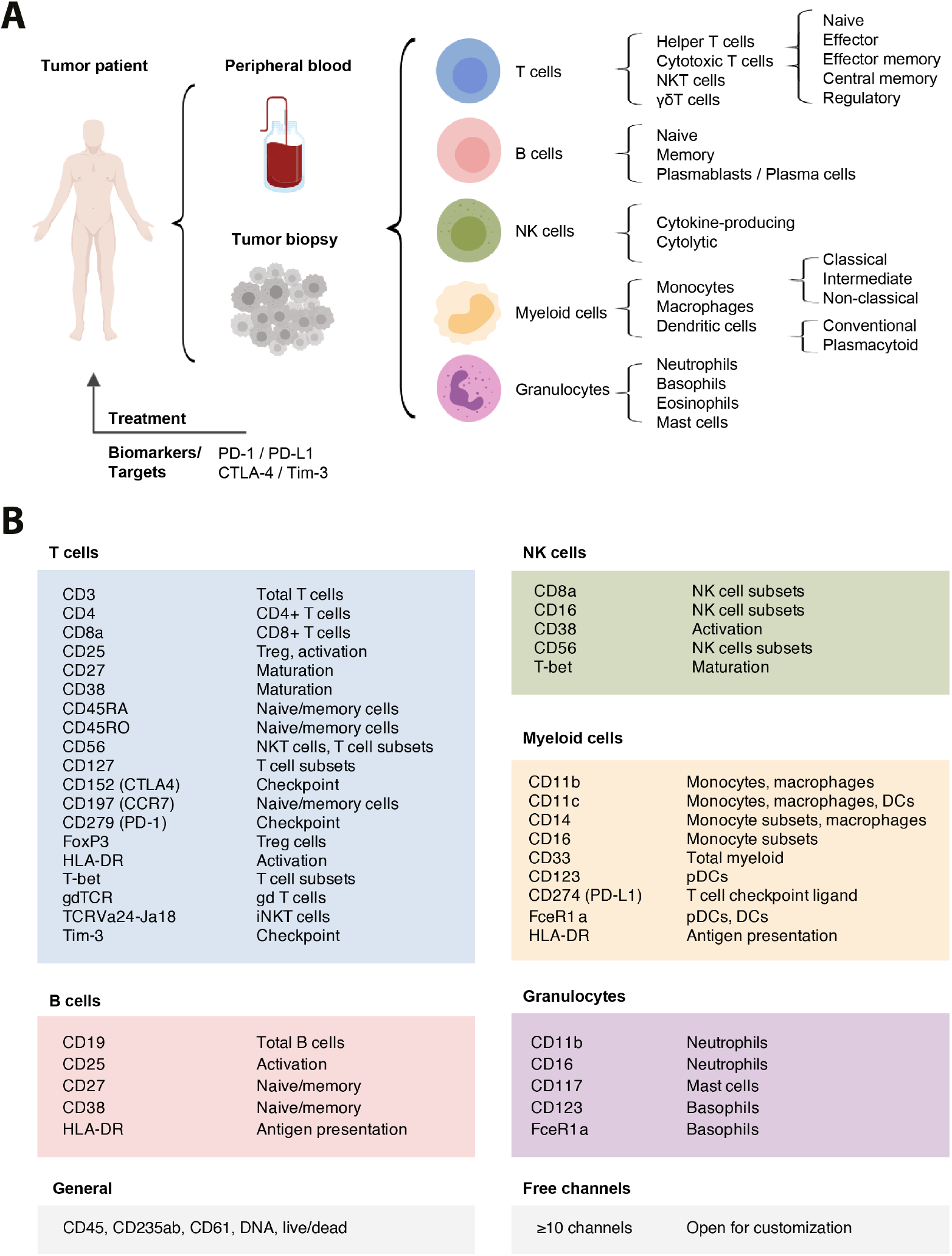
Comprehensive assessment of immune composition for clinical research in cancer immunotherapy. (A) Common sample types anticipated from tumor patients include peripheral blood samples and tumor biopsies. Within these samples, immune cell lineages and respective subpopulations are indicated. While more subsets can be delineated, these populations were chosen as a reference set of interest for comprehensive immunophenotyping. In addition to population identification, important clinical targets and currently available biomarkers are of high interest. (B) Antigens were selected based on their relevance for population and subpopulation identification or for defining important activation/maturation stages. For additional information on clones, dilutions and metal-assignments see Table S1 and the Key Resources Table.

The importance of T cells in cancer has been well-established and is illustrated by the clinical success of chimeric antigen receptor (CAR)-T cell therapies (June et al., 2018) and checkpoint blockade approaches (Ribas and Wolchok, 2018). Therefore, in addition to identifying T cells and their functionally diverse subsets, determination of the expression levels of checkpoint-related molecules such as PD-1, CTLA-4 and TIM-3, as well as receptors such as PD-L1 is critical. One specific T cell subset of high interest included in the panel is regulatory T cells (Treg). Treg cells are able to suppress T cell responses against self-antigens as well as anti-tumor T cell responses and are often associated with poor prognosis (Tanaka and Sakaguchi, 2017).

Besides T cells, functional heterogeneity also exists within other compartments, including NK cells. Traditionally, CD56^high^CD16^-^ are thought to be the main producers of an array of cytokines while CD56^low^CD16^+^ NK cells exhibit increased cytolytic activity (Björklund et al., 2016; Cooper et al., 2001). Likewise, multiple functionally diverse myeloid populations have been identified (Villani et al., 2017; Wong et al., 2012), some of which have been correlated with therapeutic success in immunotherapy (Krieg et al., 2018).

To detect and analyze the immune cell populations listed in Fig. 1A, we identified a combination of surface and intracellular proteins which characterize these immune cell lineages and their functional states (Fig. 1B) and selected a panel of 33 anti-human heavy-metal conjugated monoclonal antibodies targeting these epitopes (Table S1 and Key Resources Table). Importantly, given the high-dimensional capabilities of mass cytometry, the proposed panel does not exhaust the full range of metal isotopes commonly used in mass cytometry experiments, which allows for the inclusion of ten or more additional antibodies to further customize the panel towards more specific hypotheses. This antibody panel therefore provides the backbone needed to comprehensively and robustly identify all major immune cell populations in patient samples from immunotherapy clinical trials while allowing further customization.

### Analysis of immune composition and activation state

Having defined the range of immune cell populations and proteins to be analyzed, we utilized this panel to stain cryopreserved PBMCs from healthy donors. Stained samples were acquired on a CyTOF mass cytometer and data was normalized using bead standards (see methods). Samples were pre-gated on single, DNA^+^, live, CD45^+^ non-platelet and non-erythrocyte cells (Fig. S1A). Next, we used a sequential gating approach for initial data exploration and to identify the major immune populations within these samples (Fig. 2A). All major immune cell lineages could be readily identified using a series of lineage defining surface proteins and calculated frequencies were found to be within known ranges (Brodin and Davis, 2017) (Fig. 2B). Importantly, using the proposed gating strategy, we were able to assign 98.4 +/− 0.3% (median +/− standard error of median s.e.m.) of pre-gated cells to a specific immune lineage. Remaining cells are likely unassigned due to the strict cutoffs inherent to biaxial gating and could be identified using high-dimensional approaches as shown below.

**Figure 2.**
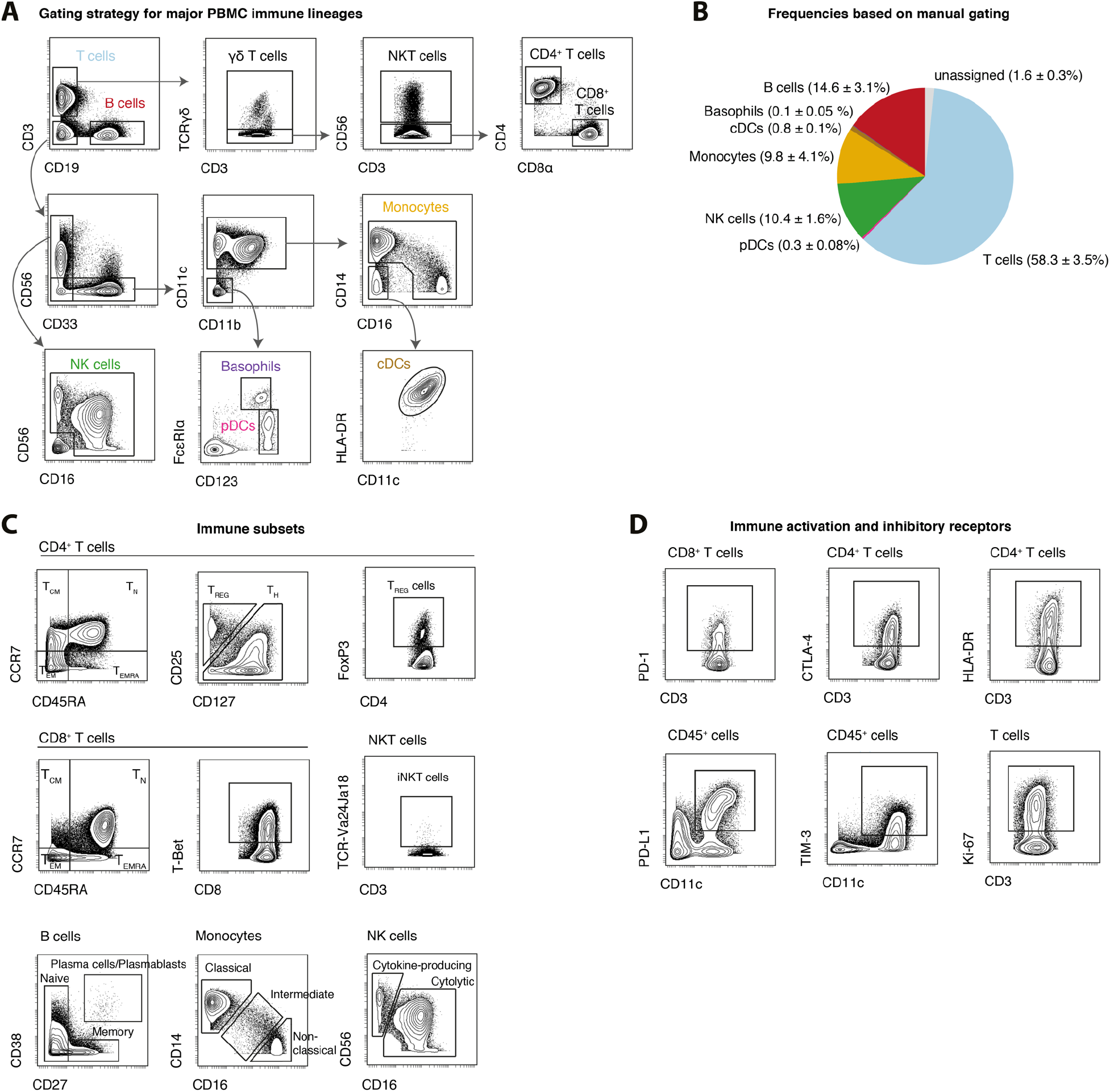
Data exploration and identification of immune cell subsets in peripheral blood. PBMCs were stained with the indicated set of antibodies (see Table S1) and analyzed by mass cytometry. (A) Cells were pre-gated as non-beads, DNA^+^, single, live, CD45^+^, CD235ab/CD61^-^, non-neutrophils (see Fig. S1). The major immune lineages and certain subsets are identified through the indicated series of gating steps. (B) Median frequencies +/− standard error of median (s.e.m.) in PBMCs from healthy donors (n = 5). (C) Exemplary identification of immune cell subsets, pre-gated on the indicated populations. Treg cells can be identified as CD25^high^ CD127^low^, FoxP3^pos^ or a combination thereof. (D) Assessment of expression levels of important checkpoint and activation molecules on various immune cell populations. Expression was induced by stimulating cells with anti-CD3, anti-CD28-coated beads for two days.

Our panel enabled the identification of multiple immune cell subpopulations. For example, T cells could be further subdivided into CD4^+^ T-helper (Th) cells, CD8^+^ T cells, natural killer T (NKT) cells and γδ T cells (Fig. 2A,C). Additionally, using differential expression patterns of CD27, CD45RA, CD45RO and CCR7, several maturation and antigen-experience states of T cells such as naïve, effector, effector memory and central memory could be discriminated (Sallusto et al., 2004) (Fig. 2C). Treg cells were identified through high expression of the interleukin-2 receptor alpha chain (CD25), low to negative levels of the IL-7 receptor CD127, and via expression of the lineage defining transcription factor FoxP3. We tested multiple staining conditions to obtain optimal intracellular staining quality for FoxP3 given its importance for Treg cell identification (Fig. S1B,C).

Aside from T cells, other immune cell lineages could be subdivided into various functional subsets (Fig. 2C). Specifically, we were able to discriminate between various stages of B cell maturation via CD27 and CD38 expression, multiple functionally distinct monocyte subsets based on their expression of CD14 and CD16, and NK cell subsets based on their combinatorial expression of CD16 and CD56.

In addition to immune cell composition, several other cellular features could be evaluated using this antibody panel (Fig. 2D). CD25, HLA-DR, and CD38 allowed determination of the activation state of T cells, while Ki-67 expression identified actively proliferating cells across multiple cell types. Importantly, expression levels of the immune checkpoint-related molecules PD-1, PD-L1, CTLA-4 and TIM-3 could be assessed on all cells. Taken together, the highly optimized approach proposed here for immune monitoring antibody allowed us to comprehensively assess both immune composition and cell activation states, simultaneously.

### Reliability and robustness across different analysis conditions

In order to assess the reliability and robustness of this immunophenotypic antibody panel in obtaining comprehensive population enumeration, we calculated the number of detected antigens on each individual cell. We found that 99.8 +/− 0.1% (median +/− s.e.m.) of live cells were positive for at least four or more antigens in our panel (Fig. 3A and Fig. S2A,B). The same was true for virtually all individual immune cell lineages, demonstrating the antibody panel’s ability to further subdivide these populations (Fig. 2B). While certain antigens might be downregulated in specific diseases, in contexts with substantial cell activation, this number will likely increase as additional proteins become expressed. Importantly, expression of a board range of proteins ensures that all major immune lineages differ from each other by expression of multiple proteins (Fig. S2C), indicating that cell identification does not depend on a single antigen given an appropriate gating strategy or by using clustering approaches as discussed below.

**Figure 3.**
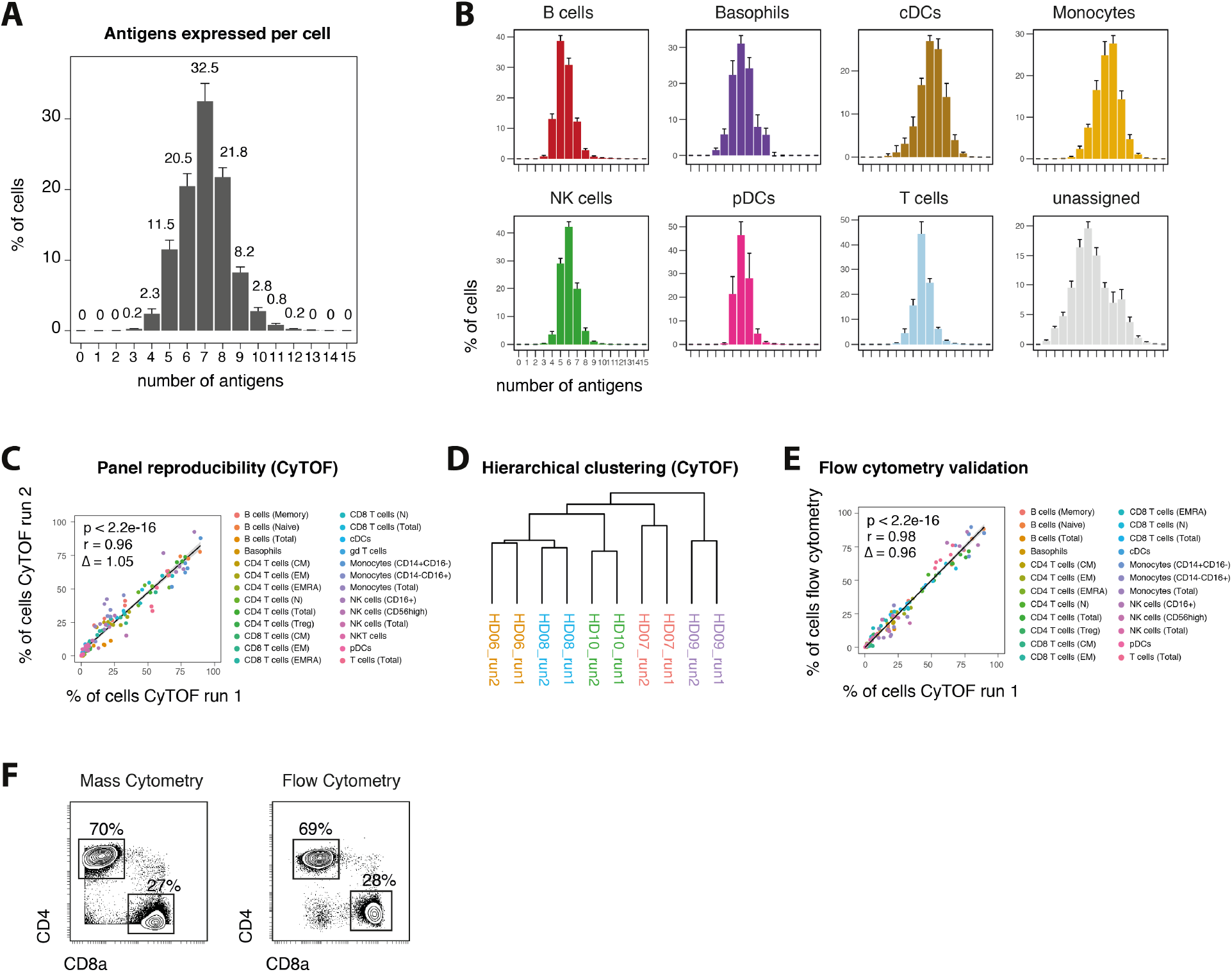
Reproducible assessments of immune composition across independent analyses. PBMCs from healthy donors (n = 5) were analyzed in multiple research centers. Immune cell populations were identified through serial gating as before (see Fig. 2). (A) Median number positive antigens per cell, based on manually determined cutoffs (see Fig. S2A). Numbers indicate median frequency of total pre-gated cells. Error bars represent s.e.m. (B) Median number of positive antigens per cell as in A, stratified by immune cell lineage. (C) Different PBMC aliquots of the same donors (n = 5) were stained and acquired by mass cytometry in two different research institutes. Frequencies of immune lineages were determined through serial gating. Linear regression line is shown in black with the 95% confidence intervals (CI, shaded). Coefficients, p-values and slope D were calculated based on data from all donors. (D) Hierarchical clustering of samples from different mass cytometry runs based on frequencies as in C. (E) PBMCs aliquots of the same donors as in C were stained and acquired by flow cytometry, employing four separate staining reactions. Frequencies of immune lineages were determined through serial gating and plotted against the frequencies determined from mass cytometry as in C. Linear regression line is shown in black with the 95% confidence intervals (CI, shaded). Coefficients, p-values and slope D were calculated based on data from all five donors. (F) Exemplary biaxial plots and frequencies of CD4^+^ and CD8^+^ T cell subsets within one donor (HD08), as determined by mass cytometry (left) and flow cytometry (right).

To assess the robustness of the selected panel across different research institutions, aliquots of PBMC samples obtained from the same blood draw of five healthy donors were distributed to two research centers where sample staining was performed by the respective researches, using separate reagents. Stained samples were then acquired on the two respective mass cytometers present in these laboratories. Immune cell frequencies were centrally determined through manual gating and compared between the individual runs (Fig. 3C). We found strong agreement (r = 0.96) between the manually gated immune cell populations from the two independent runs. This correlation was found over a broad range of frequencies and not dependent on highly abundant populations (Fig. S2D). Further, frequency-based hierarchical clustering grouped aliquots from the same donor run on different CyTOF analyzers together, thus confirming the data reproducibility between different study centers (Fig. 3D). Additional aliquots of the same PBMCs were run by flow cytometry, employing four independent antibody panels focusing on separate immune cell populations. Importantly, we obtained strong agreement (r = 0.98) between immune cell populations over a broad range of frequencies analyzed with either flow cytometry or CyTOF (Fig. 3E,F and Fig. S2D).

Lastly, to accommodate for a wide variety of immune sample collection techniques, we assessed effect of sample fixation prior to surface staining and analysis using mass cytometry (Fig. S2E-G). We calculated fold changes of the 95^th^ percentile between unfixed and PFA-fixed cells for each antigen. While the majority of antigens were not overtly altered in their dynamic ranges, a subset of antigens (including CCR7 and CD11b) showed decreased staining on previously fixed cells. However, manually gated immune cell frequencies from live-stained cells versus cells fixed with PFA prior to surface staining were nevertheless highly correlated (r = 0.94). As before, hierarchical clustering confirmed an overall highly similar immune profile between fixed and unfixed samples. Together, these data demonstrate the robustness and reproducibility of this mass cytometry-based analysis across multiple study centers and staining conditions as well as strong correlation with the historical gold standard, fluorescence-based flow cytometry.

### Data visualization and population identification using automated approaches

Thus far we used a defined sequential gating strategy to identify major immune cell populations, a method that is widely used by researchers and founded in empirical biological knowledge. However, with the increase in simultaneously acquired parameters, it is progressively infeasible to manually identify populations in highly multiplexed datasets, making computational approaches such as clustering extremely advantageous (Chester and Maecker, 2015; Mair et al., 2016; Saeys et al., 2016; Spitzer and Nolan, 2016).

To enable initial exploration, high-dimensional data is often projected into a lower dimensional space interpretable by humans using dimensionality reduction algorithms. These lower dimensional maps give an immediate overview of data structure and the presence of various populations. One method that has become increasingly popular is t-distributed Stochastic Neighbor Embedding (tSNE) (Amir et al., 2013; Maaten and Hinton, 2008). We visualized PBMC data from five healthy donors using tSNE and assigned cells to unique colors by overlaying the results of our manual gating (Fig. 4A and Fig. S3A). Manual gating and separation by tSNE appeared in high concordance, demonstrating consistent results with one another. Another hybrid approach that allows the visualization and comparison of multidimensional datasets are scaffold maps (Spitzer et al., 2015). Scaffold maps groups similar cells into clusters, which are then visualized based on their similarity with manually determined (e.g. gated) landmark nodes. We built a reference scaffold map using healthy donor PBMCs and used manual gating to define landmark nodes, which represented all major immune cell populations identified in our mass cytometry data (Fig. 4B). This method allows for comparison with other samples, such as tumor infiltrating leukocyte populations from tissue biopsies of cancer patients, which can then be mapped onto this reference map and compared through visual inspection or statistical methods (Spitzer et al., 2017) (Fig. 4C). Besides scaffold, a multitude of other high-dimensional clustering algorithms have been reported (Weber and Robinson, 2016). While many of these algorithms now have graphical user interfaces (e.g. cytofkit (Chen et al., 2016) and Cytosplore (van Unen et al., 2017)), comprehensive and reproducible analysis methods for large groups of samples remains a challenge that often requires basic familiarity with programming languages.

**Figure 4.**
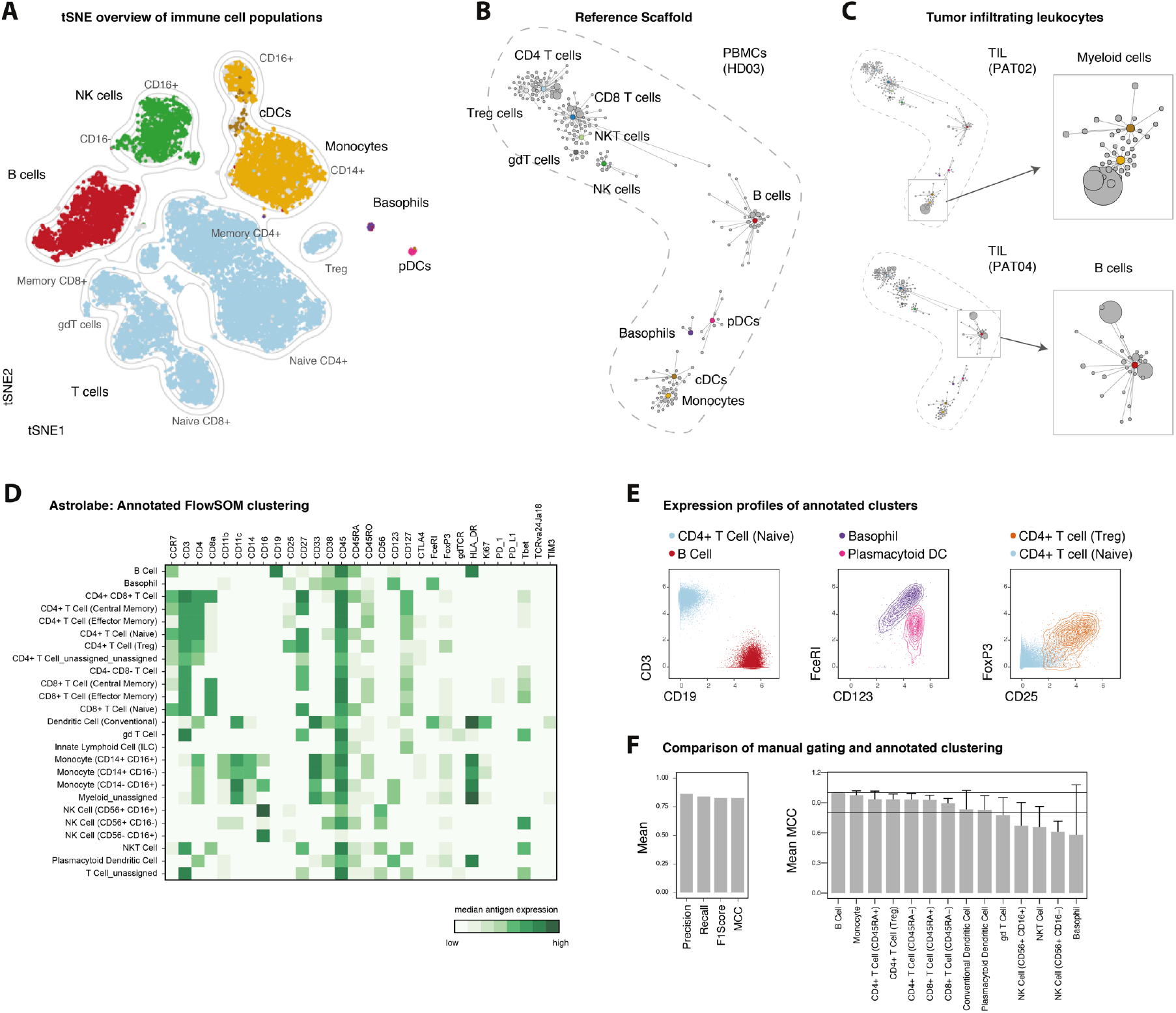
Automated data visualization and population identification. PBMCs from healthy subjects (n = 5) and tumor biopsies from cancer patients (n = 5) were analyzed by mass cytometry using the described reference panel. (A) Data from all healthy donors was randomly subsampled to 20’000 cells and subjected to tSNE dimensionality reduction. Cells are colored by their immune cell lineage assignment from manual gating. Grey indicates cells unassigned by manual gating. (B) A reference scaffold map of PBMC data was created using manually gated landmarks (colored) and all antigens for the clustering analysis. Inter-cluster connections were used to create the graph but are not depicted here. Shown is one representative sample (HD03). (C) Pre-gated, CD45^+^ cells from tumor samples were mapped onto the reference scaffold. Maps from two patients are shown (left). Enlarged examples of modulated immune cell populations are pointed out (right). (D) PBMC data as above was clustered and automatically annotated using the Astrolabe platform. Shown are median expression levels of all antigens for all clusters. (E) Exemplary expression profiles of immune cell populations as determined by Astrolabe (HD06). (F) Mean precision, recall, F1score and Matthews correlation coefficient (MCC, see methods) between manual lineage assignments and FlowSOM-based clustering for all donors and populations (left). Mean MCC for all donors stratified by population (right). Two vertical lines indicate MCC = 1 (maximum agreement) and MCC = 0.8, respectively.

Recent automated commercial solutions have been developed to analyze large multidimensional datasets. We here employed the Astrolabe platform (Astrolabe Diagnostics, Inc.) which uses the FlowSOM algorithm (Van Gassen et al., 2015) followed by a labeling step which automatically assigns cells to pre-selected and biologically known immune cell lineages (Fig. 4D,E). Depending on the required resolution, these populations can be further subdivided, again using unsupervised FlowSOM-based clustering (Fig. S3B). Using the Matthews Correlation Coefficient (MCC, see methods) to compare lineage assignments between manual gating and clustering, we found good correlations for all major leukocyte populations (Fig. 4F and Fig. S3C,D), with minor disagreements for basophils (present here at extremely low levels) and NK cell subsets (Fig. S3E).

In summary, a variety of automated methodologies can be applied to the high-dimensional data sets generated using our proposed antibody panel, thus allowing the exploration, visualization and comparison of single samples or sample groups to ultimately gain novel biological insights in a hypothesis-free and comprehensive approach.

### Identifying disease-associated immune signatures following bone marrow transplantation

One scenario in which comprehensive immunophenotyping, without prior knowledge of the system composition, is crucial is hematopoietic reconstitution in leukemia patients following bone marrow transplantation (BMT). We collected PBMC samples from fifteen individuals, sampled at multiple timepoints following BMT for a total of 28 samples (see Table S2). Of these patients, a small subset suffered from graft versus host disease (GvHD, n = 3), while most other patients did not experience such complications (Fig. 5A). In order to monitor immune reconstitution and to identify potential GvHD-associated immune signatures, we applied the above outlined mass cytometry-based experimental and analytic workflow.

**Figure 5.**
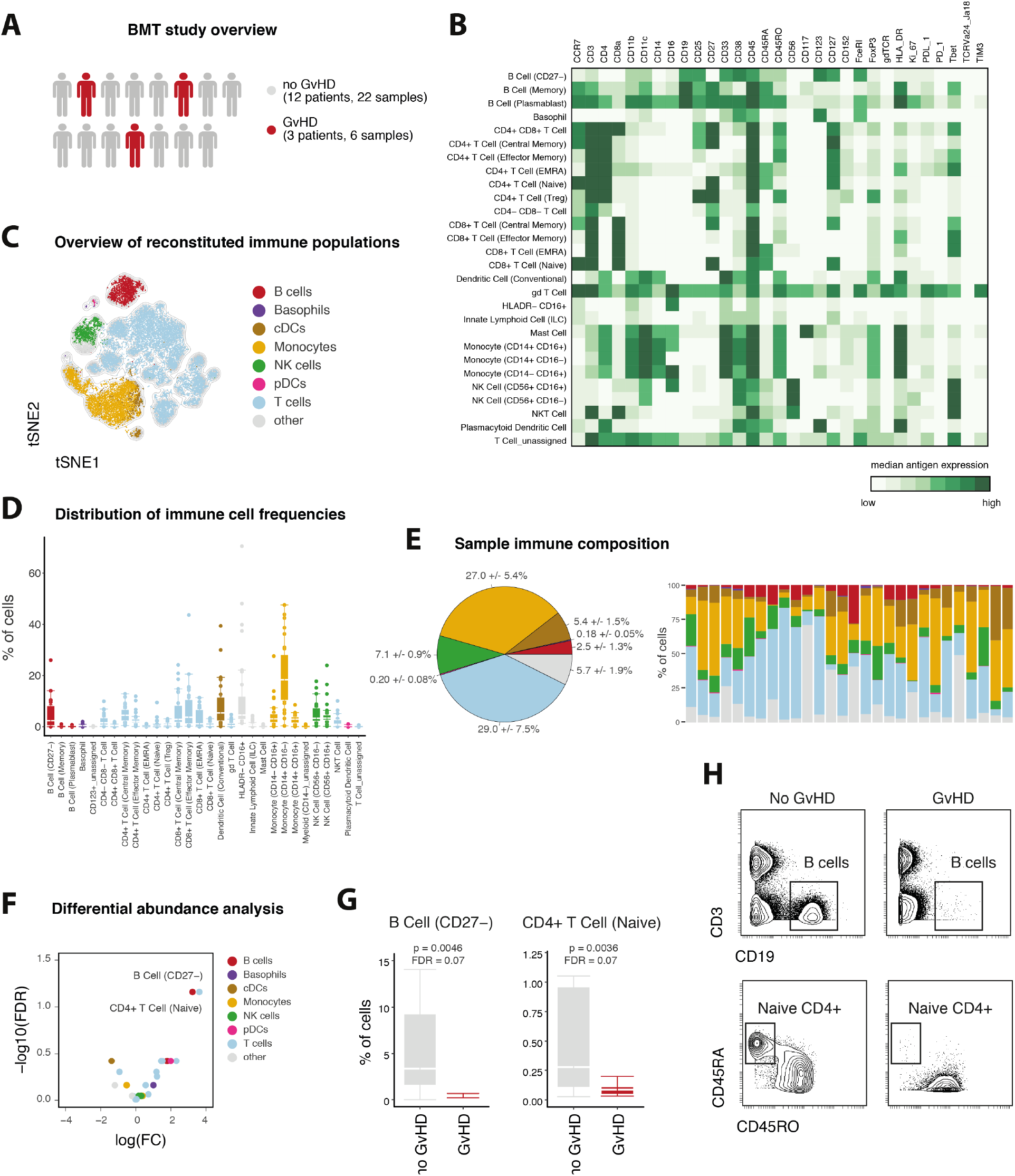
Identification of disease-associated immune signatures following bone marrow transplantation. (A) Following tumor therapy, patients (n = 15, Table S2) underwent bone marrow transplantation. Peripheral blood samples were collected and subsequently stained with the described reference panel and analyzed by mass cytometry. (B) Data was uploaded to the Astrolabe platform, clustered and automatically annotated. Exemplary heatmap of one patient depicting the median protein expression levels across all populations identified through clustering. (C) 20’000 randomly subsampled cells of one patient were subjected to tSNE dimensionality reduction. Color-assignments represent different immune lineages as identified through annotated clustering. (D) Clustering-derived frequencies of immune populations for all samples in this study (n = 28). Boxplots depict the interquartile range (IQR) with a horizontal line representing the median. Whiskers extend to the farthest data point within a maximum of 1.5x IQR. Points represent individual samples. (E) Frequencies of immune cell subpopulations were combined into frequencies for major immune cell lineages and color-coded as in C. Pie chart depicting the median frequencies +/− s.e.m. of all major immune lineages across all samples (left). Immune composition for all analyzed samples (n = 28, right). (F) False-discovery rate (FDR) and fold change (FC) of immune cell frequencies in patients with or without GvHD. (G) Comparison of differentially abundant immune cell frequencies in patients with or without GvHD. (H) Confirmation of reduced abundance of B cells (top) and naïve CD4^+^ T cells (bottom) in an exemplary patient with (right) and without GvHD (left). Examples of B cells were pre-gated on single, live, CD45^+^ cells. Examples of naïve CD4^+^ T cells were pre-gated on single, live, CD4^+^ T cells.

Following staining and acquisition, we utilized the Astrolabe platform to identify the major immune populations and their subsets. Annotated clustering identified 30 immune cell subsets spanning the major immune lineages (Fig. 5B). tSNE dimensionality reduction was used to give an immediate overview of various reconstituted populations (Fig. 5C). Across all samples, immune reconstitution was dominated by T cells (29.0 +/− 7.5%) and monocytes (27.0 +/− 5.4%), followed by NK cells (7.1 +/− 0.9%) and HLA-DR^-^CD16+ cells (Fig. 5E). B cells (dominated by CD27^-^ B cells) were present with lower frequency (2.5% +/− 1.3%).

Exploring the biological significance of patient to patient variation in their immune composition, we investigated whether immune cell proportions stratify between patients with different clinical outcomes, e.g. the occurrence of GvHD. To compare between patients with or without GvHD, we calculated fold changes (FC), p-values and false discovery rates (FDR), correcting for multiple-hypothesis testing (see methods). This approach identified a reduction in two immune cell populations as a potential immune-signature of failed engraftment/occurrence of GvHD in this cohort (Fig. 5F). Firstly, CD27^-^ B cells were reduced in patients with GvHD (0.44 +/− 0.21 vs. 3.33 +/− 2.2%, p = 0.0046, FDR = 0.069, Fig. 5G). In addition, patients with GvHD displayed lower frequencies of naïve CD4^+^ T cells (0.09 +/− 0.03% vs. 0.3 +/− 0.3%, p = 0.0036, FDR = 0.069). Lastly, while the comprehensive assessment of a broad range of immune cell populations was necessary to identify these stratifying populations, once their identity is known, manual gating can again be used to confirm their reduction in patients with GvHD (Fig. 5H). In summary, this demonstrates the utility of the outlined framework to perform clinically-relevant monitoring of immune perturbations in a medical setting. Employing this approach, treatment-, disease- or time-dependent immunological responses can be assessed in a straightforward and comprehensive manner to discover novel biomarkers and immune signatures.

### Extendibility and flexibility of the reference assay framework

Given the proposed application of this workflow to a diverse array of studies, an important feature of the immunophenotypic antibody panel is that it does not exhaust the full range of available lanthanide isotopes available for use by mass cytometry. Up to ten antibodies or more, depending on the availability of newly developed reagents, can be added to the described reference panel without modification. We illustrated this ability to customize the panel in two separate scenarios, focusing on different leukocyte populations. We targeted up to ten additional antigens with mass-tagged antibodies, stained, and acquired samples from multiple donors with these antibodies in addition to the reference panel.

First, we included an additional ten antibodies to further distinguish B cell maturation as well as co-stimulatory molecule and isotype expression (Kaminski et al., 2012) (Fig. S4A-C). Together with the immunophenotypic reference panel, these additional antigens enabled the identification of multiple additional B cell subpopulations including plasma cells and several stages of isotype switched naïve and memory B cells.

Further, we set focus on tissue-resident myeloid cell subpopulations by including antibodies against molecules associated with DCs, neutrophils, monocytes and macrophages and their activation or co-stimulatory states (Fig. 6A). For this analysis, we included tumor biopsies (n = 4) and matched, metastatic lymph node samples (n = 2) from patients with squamous cell carcinomas (see Table S2).

**Figure 6.**
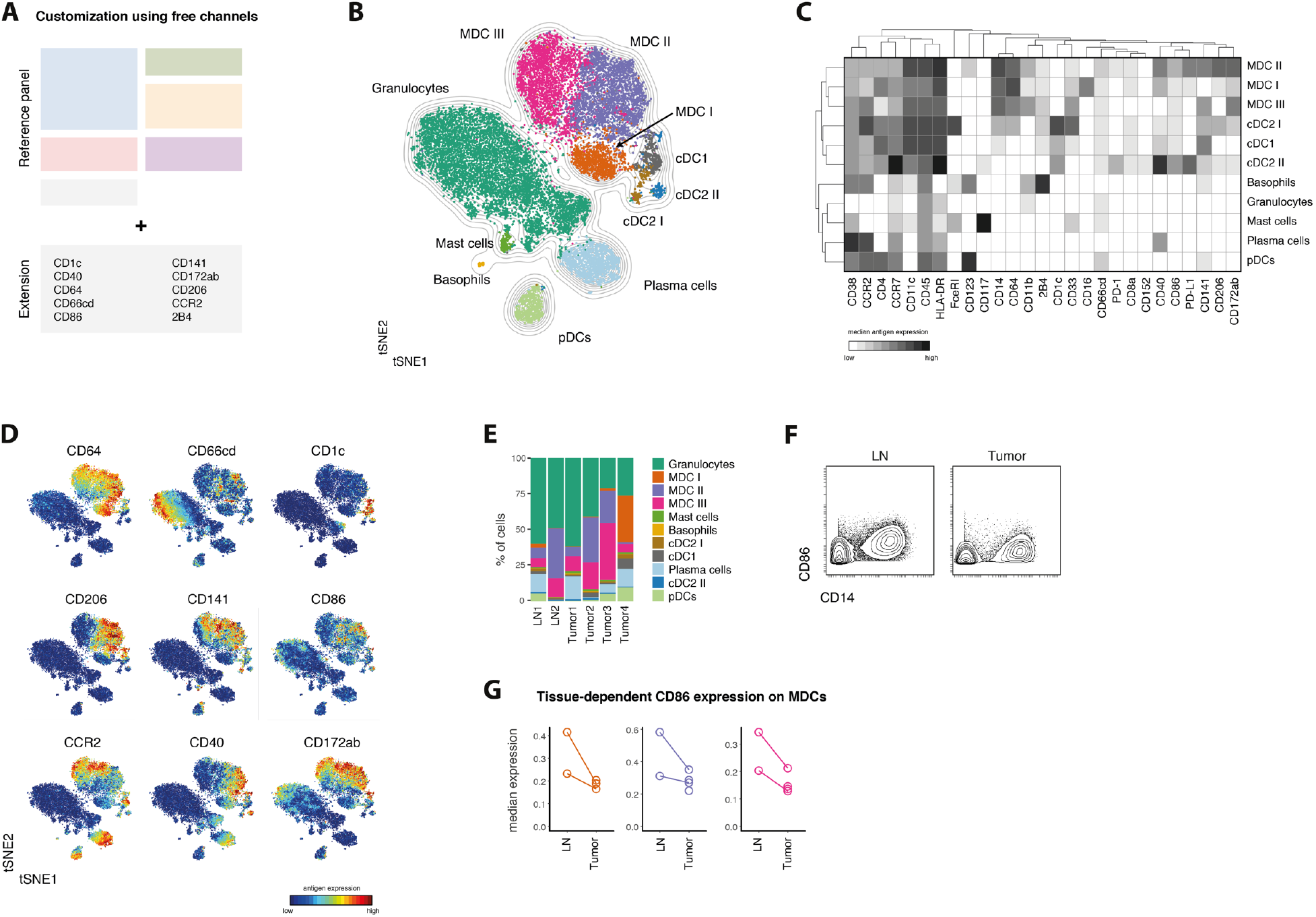
Flexibility of the proposed framework enables augmented exploration of heterogeneous populations. (A) Antibodies targeting additional antigens of interest were conjugated to non-occupied heavy metal isotopes. Cells from lymph node biopsies (n = 2) and tumor biopsies (n = 4) of patients with head and neck carcinoma (see Table S2) were stained with these antibodies in combination with the reference set. (B) Data was pre-gated on single, live, CD45^+^CD3^-^CD19^-^CD7^-^CD56^-^ to exclude T cells, most B cells and NK cells. To create a tSNE overview, data from all samples was randomly subsampled to 20’000 cells with equal contribution from all samples. Cells are colored by their FlowSOM-based cluster-assignment. Grey lines indicate the density distribution of the tSNE map. MDC: monocyte-derived cells. (C) Cluster-based median expression levels for all population relevant antigens used in the tSNE and FlowSOM analysis. (D) Protein expression levels of all additional antigens are overlaid as a color-dimension onto the tSNE map. (E) Frequencies of FlowSOM-based clusters as in B and C in all samples. (F) Exemplary CD86 expression levels on total MDCs (CD14^+^ cells) in cells derived from a lymph node metastasis (left) and primary tumor (right) of the same patient. (G) Median CD86 expression levels (arcsinh-transformed and percentile normalized) on MDC subsets from lymph nodes and tumors. Lines connect different tissues of the same patients.

Using a combination of tSNE visualization and FlowSOM-clustering, multiple myeloid subpopulations could be distinguished (Fig. 6B,C). This included previously unresolved subsets of cDCs (CD141^+^ cDC1 and CD1c^+^ cDC2) as well as different populations of monocyte/macrophage cells, hereafter referred to as monocyte-derived cells (MDCs). In addition to their identification, these subsets could also be analyzed for differential expression of many subset associated proteins (CCR2, CD244, CD172ab and CD206) and costimulatory molecules (CD40 and CD86), which have been shown reflect the activation state and propensity to provide co-stimulation to T cells (Fig. 6D).

While preliminary due to the limited number of samples, this panel extension enabled us to compare between cells isolated from primary tumors and lymph node metastases. Frequencies of defined subpopulations were comparable between tumors and lymph nodes (Fig. 6E and Fig. S4D,E). We further compared the expression of costimulatory molecules on myeloid populations from lymph nodes and tumors and observed a trend towards increased CD86 expression on all three MDC subsets isolated from lymph node metastases compared to the respective primary tumors (Fig. 6F,G). In summary, these results demonstrate that, while retaining the ability to cover all immune populations across a variety of tissues and collection conditions (Fig. 1-5), the proposed immune reference workflow provides the flexibility to further increase the resolution of the analysis towards a specific immune population or scientific hypothesis.

## Discussion

In this study, we established a reference panel of 33 anti-human antibodies for mass cytometry that can easily be incorporated into routine immunophenotyping studies in the context of cancer immunotherapy. The selected target antigens are distributed broadly across immune cell types and thus ensure that all major immune cell lineages and various functional subsets can be identified robustly and unambiguously. Apart from proteins essential for the identification of immune cell populations, we also included antibodies against targets that can be used to assess functional states, e.g. proliferative activity or expression levels of immune checkpoint-related molecules such as CTLA-4, Tim-3, PD-1 and PD-L1, some of which have already been proposed as candidate biomarkers in cancer immunotherapy (Patel and Kurzrock, 2015).

We validated the panel using various sample types from healthy donors or cancer patients, including PMBCs and biopsies of tumor tissue or lymph nodes. In all cases, we were able to identify the major immune cell lineages as well as their functionally diverse subsets and cell states. Additionally, these samples were collected and analyzed by different researchers across various research institutions, underwent different pre-processing protocols and were stained and acquired at multiple locations (Leipold et al., 2018). Notwithstanding, we obtained highly correlated results from each of the patient samples analyzed, regardless of pre-staining processing or location where the samples were analyzed. Additionally, immune cell frequencies derived from flow cytometry methods strongly correlated with our mass cytometry results (Bendall et al., 2011), further validating the proposed workflow.

It should be noted that the proposed reference panel focusses on major immune cell populations and well-established subpopulations. However, to date, there is no comprehensive consensus regarding cell type definitions and annotations, and many immune cell populations can be further subdivided depending on the use of additional antigens. To account for this and to allow customization of the antibody panel for specific research needs, the proposed immunophenotypic reference antibody panel does not exhaust the full range of available analysis channels, and additional antibodies can easily be added. Importantly, the absence of spectral overlap between different analysis channels in mass cytometry allows the straightforward addition of further antibodies. We illustrated this flexibility by including additional antibodies in the panel that were specific for B cells or myeloid cell subsets and activation states; however, other cell populations or combinations thereof could be targeted. Furthermore, fixation and permeabilization procedures are already integrated in this framework, thus allowing the incorporation of additional intracellular antibodies without having to modify the employed staining protocol. Currently, up to ten channels can be customized, and making use of alternate antibody-heavy metal conjugation protocols, such as direct binding of cisplatin to partially reduced antibodies (Mei et al., 2016), this number can be further increased. Open channels also ensure compatibility of the panel with fixed or live-cell barcoding approaches (Hartmann et al., 2018; Mei et al., 2015; Zunder et al., 2015). These approaches help to eliminate technical variability and increase sample comparability, which is especially valuable when using clinical samples obtained from different research studies. Our framework therefore further contributes to the standardization and quality control of mass cytometry experimentation, which builds upon the already published reports of using bead-based normalization protocols (Finck et al., 2013) and the addition of reference cells to increase comparability between different experiments (Kleinsteuber et al., 2016).

With minor exceptions, this panel is exclusively comprised of commercially available, off-the-shelf reagents, minimizing conjugation batch differences across time and different study sites. Implementation of such standardized experimental workflows and antibody panels have been proposed for flow cytometry (Finak et al., 2016; Maecker et al., 2012) but have not yet been implemented for analogous studies using mass cytometry. In addition to assessing a broad and defined set of immune cell populations, our proposed workflow will allow valuable cross-trial comparisons and would simplify and enhance meta-analyses as recently proposed (Hu et al., 2018). Further, we have demonstrated that results generated using this workflow are amenable to a variety of data analysis approaches. Major immune cell subsets and established subpopulations can be identified using a series of two-dimensional gates using the proposed manual gating scheme (Fig. 2). However, many alternative approaches exist and, especially for the comprehensive exploration of high-dimensional data sets, automated data analysis methods are advantageous (Newell and Cheng, 2016; Saeys et al., 2016). We utilized multiple, semi-automated algorithmic analyses approaches, including tSNE dimensionality reduction (Amir et al., 2013; Maaten and Hinton, 2008) as well as clustering and visualization through Scaffold maps (Spitzer et al., 2015). Alternatively, data could be visualized using force-directed layouts (Samusik et al., 2016) or uniform manifold approximation and projection (UMAP) dimension reduction (Becht et al., 2018; McInnes and Healy, 2018). Other approaches dedicated to identifying differential immune cell frequencies in groups of samples can be applied, including approaches relying on statistical comparisons of cluster frequencies (Bruggner et al., 2014; Spitzer et al., 2017), convolutional neutral networks (Arvaniti and Claassen, 2017), empirical Bayes moderated tests (Weber et al., 2018) or hyperspheres (Lun et al., 2017). Since these approaches typically require advanced computational skills, we additionally demonstrate compatibility of this experimental workflow with a fully-automated, commercial analysis platform (astrolabediagnostics.com) to perform a systems-level analysis of immune cell reconstitution following BMT (Lakshmikanth et al., 2017; Stern et al., 2018) and to identify factors associated with the development of acute GvHD (Stikvoort et al., 2017). Albeit preliminary due to the limited sample number and potential confounding factors, we demonstrated the utility of our framework to identify such disease-associated cellular immune signatures in a clinical cohort. Importantly, we are currently employing the described methodology to investigate the longitudinal influence of modified grafts in this scenario. In addition, this framework is already being applied to multiple studies in the field of immunotherapy research, including the study of DC vaccination approaches in combination with checkpoint inhibition (Nowicki et al., accepted).

In summary, we have established and extensively validated an experimental framework for comprehensive immunophenotyping. While the initial scope of this panel was its application to clinical research in the field of cancer immunotherapy, its broad assessment of immune cell states and populations would be a valuable approach for research in other fields such as infectious disease (Bengsch et al., 2018; Newell et al., 2012), vaccine development (Pejoski et al., 2016) and assessment of autoimmunity (Hartmann et al., 2016; Rao et al., 2017). This study demonstrates this platform’s broad applicability and provides examples of how it will accelerate and improve immune monitoring of patients enrolled in clinical trials. Altogether, by taking our cell agnostic approach to immune monitoring, laying out a unified protocol and panel for comprehensive analysis, this study democratizes the elucidation of therapeutic mechanisms and discovery of immune cell signatures and biomarkers.

## Acknowledgements

We thank D.R. Glass, D. Mrdjen and J.P. Oliviera for discussions and comments. F.J.H. was supported by the EMBO organization (EMBO Long-Term Fellowship), the Novartis Foundation for medical-biological Research and the Swiss National Science Foundation (SNF Early Postdoc Mobility). K.B.J. is supported by KL2-TR001870. M.H.S. was supported by the Chan Zuckerberg Biohub, the NIH S10 1S10OD018040-01 and DP5OD023056. S.C.B. was supported by the Damon Runyon Cancer Research Foundation DRG-2017-09 and the NIH 1DP2OD022550-01, 1R01AG056287–01, 1R01AG057915-01, 1-R00-GM104148-01, 1U24CA224309-01, 5U19AI116484-02, U19 AI104209, The Bill and Melinda Gates Foundation, and a Translational Research Award from the Stanford Cancer Institute. The Parker Institute for Cancer Immunotherapy provided core funding for this study to M.H.S. and S.C.B.

## Author Contributions

Conceptualization, F.J.H., M.H.S., S.C.B.

Methodology, F.J.H., B.S.

Validation, F.J.H., J.B., N.S., PF.G., B.S., P.K., E.O.

Formal Analysis, F.J.H. and E.D.A.

Investigation, F.J.H and J.B.

Resources, D.M, K.B.J., H.T.M, M.H.S., S.C.B

Writing – Original Draft, F.J.H.

Writing – Review & Editing, F.J.H., J.B., PF.G., K.B.J., M.H.S, S.C.B.

Visualization, F.J.H and E.D.A.

Supervision, M.H.S. and S.C.B.

Project Administration, M.H.S. and S.C.B.

Funding Acquisition, F.J.H., M.H.S., S.C.B.

## METHODS

### CONTACT FOR REAGENT AND RESOURCE SHARING

Further information and requests for resources and reagents should be directed to and will be fulfilled by the Lead Contact, Sean C. Bendall.

### EXPERIMENTAL MODEL AND SUBJECT DETAILS

#### Human subjects

##### PBMC samples

All samples from human subjects (see Table S2) were obtained and experimental procedures were carried out in accordance with the guidelines of the Stanford Institutional Review Board (IRB). Written informed consent was obtained from all subjects. For healthy donors, fresh whole human blood in heparin collection tubes or leukoreduction system (LSR) chamber contents (Terumo BCT) were obtained via the Stanford Blood Center. Samples from BMT patients were drawn on 30 and 90 post BMT. PBMCs were isolated via Ficoll (GE Healthcare) density gradient centrifugation, resuspended in fetal bovine serum (FBS, Omega Scientific) supplemented with 10% DMSO (Sigma) and stored in liquid nitrogen.

##### Tissue samples

Tissue samples (see Table S2) were collected fresh shortly after surgery and transported for processing on ice in transport medium (Leibovitz’s L-15 medium supplemented with 6 g/L glucose and 15 mM HEPES buffer). Tumor samples were then finely minced and placed into tumor dissociation buffer (transport medium, 2% fetal bovine serum (FBS), 5mg/ml collagenase IV, 0.1 mg/ml DNase I) for 45 min at 37 °C with gentle rotation. Following dissociation, cells were filtered through a 70 μm filter, centrifuged at 500 g for 5 min at 4 °C, and resuspended in PBS with 5 mM EDTA. Cells were then mixed with viability buffer (PBS, 5 mM EDTA, 50 μM cisplatin) for 60 seconds at room temperature, quenched with wash buffer (PBS, 5 mM EDTA, 0.5 % bovine serum albumin), centrifuged at 500 g for 5 minutes at 4°C, resuspended again in wash buffer, and fixed in 1.6% PFA for 10 min at RT. After fixation, cells were centrifuged at 600 g for 5 min at 4 °C, rinsed with wash buffer, centrifuged again at 600 g for 5 min at 4°C, resuspended in freezing medium (PBS, 10% DMSO, 0.5% bovine serum albumin), and frozen at −80 °C until staining.

### METHOD DETAILS

#### Heavy-metal conjugation of antibodies

Most antibodies were obtained pre-conjugated to heavy-metal isotopes from Fluidigm. Where needed, in-house conjugations were performed using the MaxPar X8 antibody-labelling kit (Fluidigm). In short, antibody buffer exchange was performed by washing 100 μg of antibody with R buffer (Fluidigm) using a 50 kDa MWCO microfilter (Millipore) and centrifuging for 10 min, 12’000 g at RT. Antibodies were then reduced with 100 μl of 4 mM TCEP (Thermo Fisher) for 30 min at 37 °C and washed two times with C buffer (Fluidigm). Metal chelation was performed by adding lanthanide metal solutions (final 0.05 M) to MaxPar chelating polymers in L-buffer (both Fluidigm) and incubating for 40 min at RT. Metal-loaded polymers were washed twice with L-buffer using a 3 kDa MWCO microfilter (Millipore) by centrifuging for 30 min, 12’000 g at RT. Partially reduced antibodies and metal-loaded polymers were incubated together for 60-120 min at 37 °C. Conjugated antibodies were washed four times with 400 μl W buffer (Fluidigm) and collected by two centrifugations (2 min, 1’000 g, RT) with 50 μl of W buffer into an inverted column in a fresh 1.6 ml collection tube. Protein content was assessed by NanoDrop (Thermo Fisher) measurement, antibody stabilization buffer (Candor Bioscience) was added to a final volume of at least 50 v/v % and antibodies were stored at 4 °C.

#### Mass cytometry workflow

##### Sample preparation

Cryopreserved PBMC and tumor biopsy samples where thawed into 10 ml of cold cell culture medium (RPMI-1640 (life technologies), 10% FBS, 1x L-glutamine, 1x penicillin/streptomycin (Thermo Fisher)) supplemented with 20 U/ml sodium heparin and 0.025 U/ml benzonase (Sigma) and washed once (250 g, 4 °C).

##### Cellular barcoding

Where indicated, samples where barcoded and combined into a composite sample before surface staining. Barcoding was performed employing either a palladium-based barcoding approach applicable to fixed cells (Zunder et al., 2015) or a live cell barcoding methodology involving antibodies against the surface molecules beta-2-microglobulin and a sodium-potassium pump (CD298) as described (Hartmann et al., 2018).

##### Viability staining

Cisplatin (Sigma) was resuspended to 100mM in DMSO, pre-conditioned for 48 h at 37 °C and stored at −20 °C. Viability staining was performed by resuspending the sample in 1 ml of PBS and adding cisplatin to a final concentration of 500 nM, followed by incubation for 5 min at RT and washing with CSM. Where indicated, cells were fixed with 1.6% PFA in PBS for 10 min at RT and washed twice with cell staining medium (CSM: PBS with 0.5 % BSA and 0.02 % sodium azide (all Sigma)) before staining. In case live cell barcoding was employed, viability assessment was performed by substituting cisplatin with DCED-palladium (Sigma) and following the protocol as described here.

##### Antibody staining

Cell-surface antibody master-mix (2x) was prepared by adding appropriate dilutions of all cell-surface antibodies (Table S1 and Key Resources Table) into 50 μl CSM per sample. If samples contained more than 3 × 10^6^ cells, antibody volume (but not total CSM volume) was increased accordingly (e.g. 2-fold for up to 6 × 10^6^ cells). The antibody master-mix was then filtered through a pre-wetted 0.1 μm spin-column (Millipore) to remove antibody aggregates and 50 μl were added to the sample resuspended in 50 μl of CSM. After incubation for 30 min at RT, cells were washed once with CSM. For intracellular staining, cells were fixed using the FoxP3 / transcription factor staining buffer set (Thermo Fisher Scientific) to fix for 1 h at RT. After fixation, samples were washed once with CSM and once with 1x permeabilization buffer (Thermo Fisher Scientific) by centrifugation for 5 min, 600 g at 4 °C. Intracellular antibody master-mix (2x) was prepared analogously to the surface antibody mix by adding appropriate dilutions of all intracellular antibodies (see Table S1 and Key Resources Table) into 50 ul permeabilization buffer per sample. 50 μl of 2x antibody master mix was added to the samples in 50 μl permeabilization buffer and incubated for 1 h at RT. Cells were washed once with permeabilization buffer and once with CSM. Finally, samples were resuspended in intercalation solution (1.6% PFA in PBS and 0.5 μM iridium-intercalator (Fluidigm)) for 20 min at RT or overnight at 4 °C.

##### Data acquisition

Before acquisition, samples were washed once in CSM and twice in ddH_2_O and filtered through a cell strainer (Falcon). Cells were then resuspended at 1 × 10^6^ cells/mL in ddH_2_O supplemented with 1x EQ four element calibration beads (Fluidigm) and acquired on a CyTOF2 mass cytometer (Fluidigm).

#### Flow cytometry

PBMC samples were thawed as described above and subsequently treated with Fc blocking reagent (BioLegend) for 10 min at 4 °C. Antibody cocktails were then added for 30 min and incubated at 4 °C. All samples were washed with PBS containing BSA (0.5%), then fixed with 1.6% PFA for 10 min at RT. Finally, the samples were washed and analyzed on an LSRII flow cytometer (BD Biosciences) equipped with 405, 488, 561, and 640nm lasers.

### QUANTIFICATION AND STATISTICAL ANALYSIS

#### Data normalization and gating

After acquisition, data from acquired samples was bead-normalized using matlab-based software (Finck et al., 2013). Barcoded cells were assigned back to their initial samples using matlab-based debarcoding software (Zunder et al., 2015). Normalized data was then uploaded onto the Cytobank analysis platform (Kotecha et al., 2010) to perform initial gating and population identification using the indicated gating schemes (Fig. 2 and supplemental Fig. S1).

#### Data visualization and analysis

For further downstream analysis, pre-gated data was imported into the R environment (R Development Core Team, 2008) using the flowCore package (Ellis et al., 2009). Data was transformed with an inverse hyperbolic sine (arcsinh) transformation using a cofactor of 5 and normalized to the 99.5^th^ percentile of each respective channel before downstream tSNE and Scaffold analysis. Visualization of samples by tSNE dimensionality reduction was calculated using the Rtnse package (Krijthe, 2015) with default parameters: perplexity=30, theta=0.5, max_iter=1000 using the indicated channels.

To build a reference scaffold, bead and percentile-normalized data from live, CD45^+^, single, non-neutrophil cells was imported into the statisticalScaffold package (Spitzer et al., 2017). All available channels were used to build the reference maps. All population-relevant antigens were included in the clustering analysis. Astrolabe analysis was carried out by uploading bead-normalized data. Single-cell data was clustered using the FlowSOM R package (Van Gassen et al., 2015). Cell subset definitions follow (Finak et al., 2016; Maecker et al., 2012). Cluster labeling, method implementation, and visualization were done through the Astrolabe Cytometry Platform (Astrolabe Diagnostics, Inc.).

#### Statistical analysis

Cell frequencies are reported as medians unless stated otherwise. Standard error of median was calculated in R using bootstrapping with 1000-fold resampling. For frequency correlations between different centers and technologies, manually gated frequencies of cell populations were compared by linear regression using the lm() function. Hierarchical clustering using the R function hclust() was performed using the same frequency matrix.

To compare manual gating with automated clustering we employed the Matthews correlation coefficient (MCC) (Boughorbel et al., 2017; Matthews, 1975) which takes into account true and false positives as well as negatives and expresses these results in a single coefficient. A coefficient of +1 represents perfect agreement.

Differential abundance analysis for identifying GvHD-associated immune signatures was done through the Astrolabe platform using the edgeR R package (McCarthy et al., 2012; Robinson et al., 2010) following the method outlined in (Lun et al., 2017). Samples from both timepoints were pooled for this analysis.

#### Visualization

Plots were created using the ggplot2 R package (Wickham, 2016). Schematic representations were created with biorender (https://biorender.io/). Figures were prepared in Illustrator (Adobe).

### DATA AND SOFTWARE AVAILABILITY

Link to data: [data will be made available on flowrepository.org]

## Supplemental Information

### Supplementary Tables

Table S1. Reference panel of anti-human antibodies for mass cytometry.

Table S2. Donor characteristics.

Table S3. Extension panels.

### Supplementary Figures

Figure S1. Identification of single, live leukocytes and optimization of FoxP3 staining.

Figure S2. Robustness of immune cell lineage identification.

Figure S3. Automated profiling of immune cell populations.

Figure S4. Exploration of heterogeneous populations through panel extensions.

